# Cerebellar Normative Modeling Identifies Neuroanatomical Biotypes Predicting dTMS Response in Spinocerebellar Ataxia Type 3

**DOI:** 10.64898/2026.06.24.734389

**Authors:** Kaikai Wang, Yuwei Hu, Xingang Wang, Congying Chu, Lingzhong Fan, Chen Liu

**Author notes:** These authors contributed equally to this work (Kaikai Wang, Yuwei Hu). **Corresponding Authors:** Chen Liu. and Lingzhong Fan.

## Abstract

**Background:** Spinocerebellar ataxia type 3 (SCA3) presents with significant clinical heterogeneity. Traditional case-control neuroimaging, based on group means, obscures inter-individual anatomical variability, hindering the identification of stratification biomarkers for interventions like Transcranial Magnetic Stimulation (TMS).

**Methods:** To quantify individual neuroanatomical deviations, we constructed a cerebellar normative model using a multi-center dataset of 2,071 healthy controls with 2,549 MRI scans. Gray matter volume deviations (Z-scores) were mapped across 27 cerebellar lobules in 114 genetically confirmed SCA3 patients, and unsupervised clustering was applied to identify neuroanatomical biotypes. Clinical relevance was assessed by associating biotypes with ataxia severity and deep TMS (dTMS) outcomes in a longitudinal subset .

**Results:** We identified two distinct biotypes: Biotype 1 exhibited relative structural preservation (positive deviations) predominantly in the posterior cerebellum (lobules VIIB, VIIIA), whereas Biotype 2 was characterized by extensive atrophy (negative deviations) centered on the anterior motor cerebellum (lobules I-VI). Clinically, Biotype 2 patients presented with significantly more severe baseline ataxia. However, regarding treatment response, an inverse relationship was observed: Biotype 2 patients demonstrated significantly greater symptomatic improvement following dTMS compared to Biotype 1. To further identify the optimal neuromodulatory strategy for each biotype, we compared the therapeutic efficacy of repetitive TMS (rTMS) and dTMS. While both biotypes showed clinical improvement following rTMS, Biotype 1 exhibited a superior therapeutic response to rTMS relative to dTMS. Furthermore, feature weight analysis identified atrophy of the right lobule VIIB as a critical predictor of clinical severity in Biotype 2.

**Conclusion:** This study demonstrates that normative modeling can decode SCA3 heterogeneity. The identification of these biotypes reveals a dissociation between baseline structural integrity and neuromodulatory responsiveness, suggesting that patients with severe anterior cerebellar atrophy may, counterintuitively, derive greater therapeutic benefit from dTMS. Furthermore, by comparing the therapeutic efficacy of rTMS and dTMS, we further clarified biotype-specific treatment responses. These findings support the use of individualized neuroanatomical mapping for patient stratification in precision medicine.

## Introduction

Spinocerebellar ataxia type 3 (SCA3), caused by a CAG repeat expansion in the *ATXN3* gene, is the most prevalent autosomal dominant ataxia globally.^1,2^ The disease follows an inexorable progressive course, imposing a heavy burden of motor disability and non-motor dysfunction.^3–5^ Despite elucidating the genetic trigger, disease-modifying pharmacological therapies remain elusive, and current management is largely symptomatic and supportive.^6,7^ In this therapeutic void, non-invasive brain stimulation, particularly Transcranial Magnetic Stimulation (TMS), has emerged as a promising avenue to alleviate cerebellar ataxia by modulating neural excitability and plasticity.^8–10^ However, clinical translation is hindered by a critical bottleneck: therapeutic response is highly variable, with some patients showing dramatic improvement while others remain refractory. This unpredictability likely stems from the unrecognized biological heterogeneity among patients, underscoring an urgent need for biomarkers that can stratify patients and predict therapeutic responsiveness. Compounding this challenge, the optimal TMS protocol for cerebellar ataxia remains unsettled. Repetitive TMS (rTMS) has shown preliminary efficacy^11^.Deep TMS (dTMS) has been explored to augment the therapeutic benefits of neuromodulation in ataxia, while these advanced protocols hold theoretical appeal, head-to-head comparisons establishing their relative efficacy are lacking, and it remains unknown whether specific patient subgroups might preferentially benefit from one approach over the other. This therapeutic ambiguity—whether to employ rTMS or dTMS, and for whom—represents a critical unresolved question that impedes evidence-based clinical decision-making.

To resolve this heterogeneity, identifying robust neuroanatomical substrates is essential. While SCA3 pathology extends to the brainstem and basal ganglia, the cerebellum remains the indisputable epicenter of neurodegeneration.^12–21^ Post-mortem studies consistently pinpoint Purkinje cell loss as the hallmark pathology, and in vivo imaging confirms that cerebellar gray matter atrophy is the strongest structural correlate of ataxia severity.^22,23^ Thus, the cerebellum serves as the ideal “structural window” to decipher the biological variability underlying diverse clinical phenotypes and treatment outcomes.

However, mapping these alterations with precision to guide individual treatment remains a challenge. Traditional case-control studies relying on group-level averages inherently treat SCA3 as a homogeneous entity, masking subtle, idiosyncratic deviation patterns. To overcome this limitation, normative modeling has gained recognition in neuroimaging as a transformative framework for precision medicine.^24–28^ Analogous to pediatric growth charts, this approach allows for the quantification of individual deviations (Z-scores) from a healthy lifespan trajectory, effectively disentangling disease-specific pathology from normal age-related variations.^29^ While this framework has successfully mapped heterogeneity in psychiatric and neurodevelopmental disorders,^30–32^ and more recently in cerebellar pathology related to mental illness,^33–35^ its potential to decode the structural heterogeneity of hereditary ataxias like SCA3 remains unexplored. By applying this established framework to the cerebellum, we can transform complex anatomical data into standardized “deviation maps” for each patient. This shift from “average atrophy” to “individual deviation” is pivotal: it enables the use of unsupervised clustering to reveal latent neuroanatomical biotypes—subgroups with distinct spatial patterns of damage that may harbor unique potentials for neural compensation and TMS response.

In the present study, using a normative reference model established from a large multi-center healthy dataset as a framework, we systematically investigated the neuroanatomical heterogeneity in patients with SCA3. We aimed to: (1) quantify individualized cerebellar structural deviations; (2) identify distinct neurobiological biotypes based on these deviation patterns; and (3) validate the clinical utility of these biotypes, specifically testing their capacity to predict TMS treatment outcomes and to establish evidence-based recommendations for personalized TMS modality selection in clinical practice. We hypothesized that SCA3 comprises distinct neuroanatomical biotypes reflecting different compensatory capacities (e.g., structural preservation vs. exhaustion), and that these biotypes possess differential sensitivity to neuromodulation, thereby providing a long-awaited biomarker for stratified therapy and precision medicine.

## Materials and Methods

### Participants and Study Design

The study cohort comprised two primary datasets aimed at constructing a robust normative model and evaluating pathological deviations in SCA3. Dataset 1:To model the healthy lifespan trajectory of cerebellar gray matter volume, we aggregated high-resolution T1-weighted MRI scans from three large-scale public repositories: the Human Connectome Project Young Adults (HCP-YA, aged 22–37 years),^36^ the Human Connectome Project Aging (HCP-Aging, aged 36–90 years),^37^ and healthy controls from the Alzheimer’s Disease Neuroimaging Initiative (ADNI, aged 56–93 years).^38,39^ Dataset 2: SCA3 patients and healthy controls were recruited from Southwest Hospital, Army Medical University (Chongqing, China), with peripheral blood collected specifically from SCA3 patients to determine the CAG repeat length in exon 10 of the ATXN3 gene coding sequence, using polymerase chain reaction (PCR) and capillary gel electrophoresis. In Dataset 2, 40 patients received dTMS and 25 received rTMS, with two entirely independent cohorts assigned to each stimulation modality, and corresponding treatment outcomes were recorded. Detailed parameters are provided in the Supplementary Materials.

Following stringent quality control (Supplementary Fig. 1), the final analyses included 2,071 healthy controls (HCs, aged 19–93 years,a total of 2,549 scans) and 114 genetically confirmed SCA3 patients (aged 21–70 years) (Table 1). All structural MRI data were acquired using 3.0-T scanners. Detailed acquisition parameters for each site are provided in the Supplementary Materials. The study protocol was approved by the local ethics committees, and written informed consent was obtained from all participants.

**Table 1.** Demographic characteristics of the normative reference cohort and the SCA3 validation cohort.

| Cohort / Dataset | N | Age (years) | Sex(Male/Female) | Scanner Field Strength |
| --- | --- | --- | --- | --- |
| Normative Reference Cohort <sup>a</sup> | 2,071 | - | 914 / 1,157 | - |
| HCP-YA | 1,033 | 28.83 ±3.71 | 444 / 589 | 3.0 T |
| HCP-Aging | 636 | 59.57 ±14.76 | 280 / 356 | 3.0 T |
| ADNI | 153 | 75.20 ± 6.91 | 77 / 76 | 3.0 T |
| SWH-HC (Local Controls) | 249 | 51.06 ± 12.33 | 113 / 136 | 3.0 T |
| SCA3 Validation Cohort | 114 | 42.13 ± 11.52 | 64 / 50 | 3.0 T |
| Statistical Comparison <sup>b</sup> |  |  |  |  |
| Test Statistic | - | U = 113381.00 | $\chi^2 = 5.82$ | - |
| P-value | - | ns | < 0.05 | - |

Footnotes: Data are presented as mean ± standard deviation (SD) or number (N). Abbreviations: HCP, Human Connectome Project; ADNI, Alzheimer's Disease Neuroimaging Initiative; SWH, Southwest Hospital.
<sup>a</sup>The normative reference cohort integrates high-resolution T1-weighted MRI data from four sources to cover the adult lifespan
<sup>b</sup>Statistical comparisons were performed between the normative reference cohort and the SCA3 validation cohort.
The $\chi^2$ value for sex distribution was obtained by the Chi-square test
The U values were obtained by the Mann-Whitney U test
ns,not significant ( $p > 0.05$ )

### Clinical Assessment

The severity of ataxia and daily functional disability were evaluated using the International Cooperative Ataxia Rating Scale (ICARS)^40^ and the Scale for Assessment and Rating of Ataxia (SARA).^41^

### Image Processing and Cerebellar Parcellation

Cerebellar segmentation was performed using the ACAPULCO algorithm,^42^ a state-of-the-art deep learning framework integrated into the ENIGMA cerebellar pipeline. Leveraging a cascaded dual-convolutional neural network (CNN) architecture, ACAPULCO provides rapid, high-fidelity parcellation of cerebellar lobules.^43^ Before segmentation, image preprocessing involved two key steps: (1) N4 bias field correction to rectify intensity inhomogeneity^44^; and (2) rigid registration to the MNI-space ICBM 2009c template using Advanced Normalization Tools (ANTs) for standardized spatial alignment.^45^ Following segmentation, gray matter volumes (mm³) were extracted for 27 regions of interest (ROIs), comprising: bilateral lobules I–VI, Crus I, Crus II, VIIB, VIIIA, VIIIB, IX, and X, as well as vermian lobules VI–X. Intracranial volume (ICV) was estimated for all participants using FreeSurfer (surfer.nmr.mgh.harvard.edu) to account for global head size variations.

### Normative Modeling of Cerebellar Gray Matter Volume

The normative model was implemented using the PCNtoolkit (https://pcnportal.dccn.nl/)^25,46^ in Python 3.9.

#### Data Partitioning

To ensure objective benchmarking and prevent overfitting, the aggregate HC cohort was randomly partitioned into a training set (70%) and an independent test set (30%), stratified by age, sex, and scanning site. Rationale: It is crucial to note that the model was trained exclusively on the 70% HC subset. This design mitigates potential bias; including the test HCs in training could artificially inflate the model’s fit for controls relative to patients, potentially leading to false-positive deviation signals in the SCA3 group.^33,47^

#### Modeling Strategy

We employed Bayesian Linear Regression (BLR) combined with likelihood warping to model non-Gaussian distributions.^48^ Predictors included age, sex, and ICV. Crucially, the scanning site was modeled as a fixed effect to rigorously correct for multi-center batch effects,^49,50^ a method proven to yield robust site-invariant predictions.^33,49^

#### Deviation Quantification

For each subject (*i*) and cerebellar lobule (*j*), the individual deviation was quantified as a Z-score:

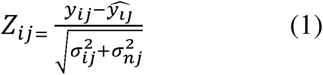

where *y_ij_* is the observed gray matter volume, *ŷ_ij_* is the predicted mean volume, σ^2^_*ij*_ represents predictive uncertainty, and σ^2^_*nj*_ denotes model noise variance.^48^ Model performance was evaluated on the independent HC test set using explained variance (EV) and mean squared log-loss(MSLL).^51^ Extreme deviations were defined as |*Z*| > 1.96 (95% confidence interval).^33,34^

### Assessment of Individualized Deviations in SCA3

Individualized deviation maps for SCA3 patients were generated by projecting their raw volumetric data onto the normative reference model (Figure 1B). To characterize disease-specific pathology, we compared the frequency and magnitude of extreme deviations between SCA3 patients and HCs using the two-sample t test or the Mann–Whitney U test, where appropriate. We further generated spatial overlap maps to visualize the heterogeneity of anatomical damage across the patient cohort.

**Figure 1.**
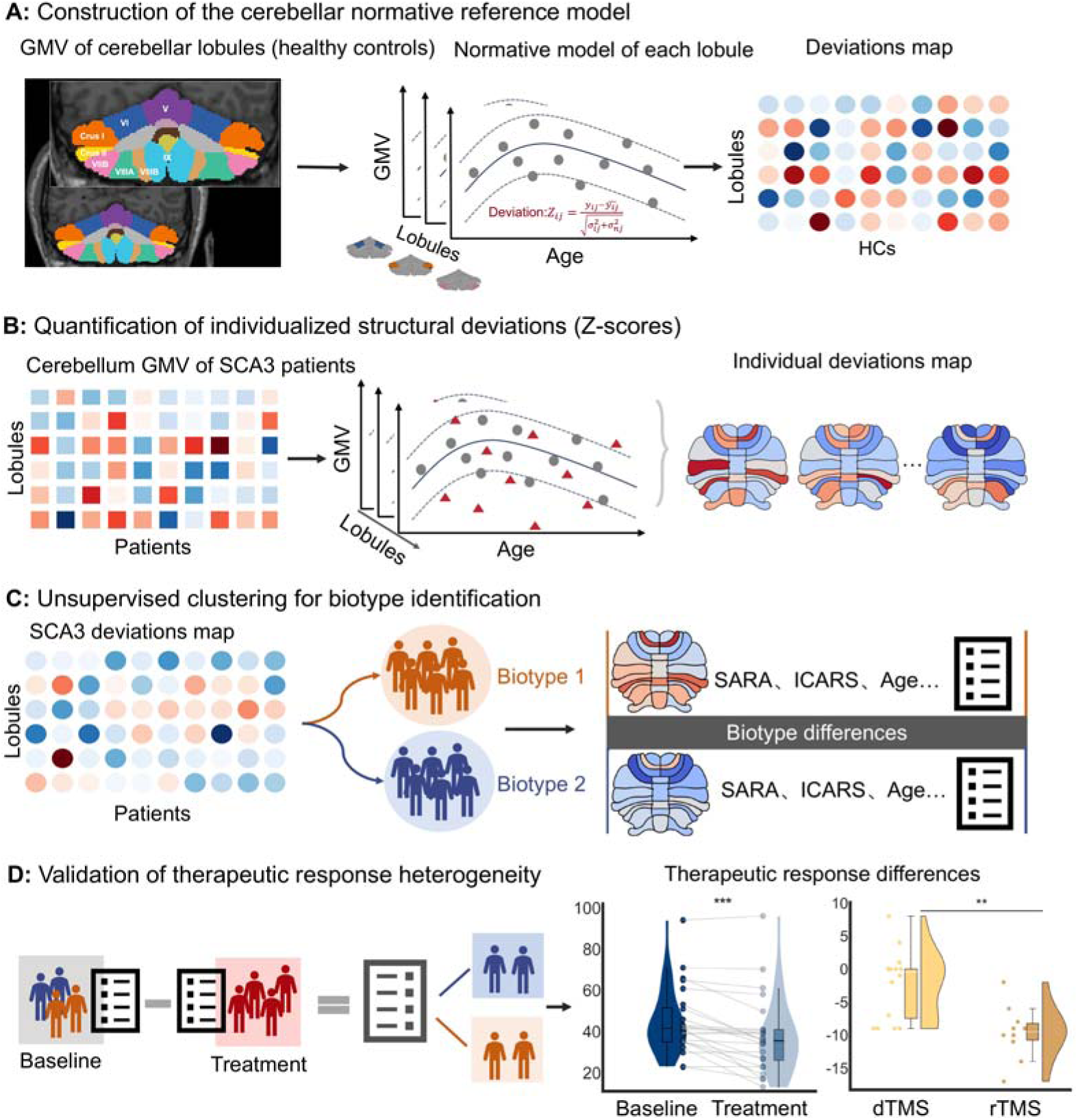
Study workflow for normative modeling and biotype identification. **(A)** Construction of the cerebellar normative model using Bayesian Linear Regression on a training set of healthy controls (HCs, gray dots) to map age-related gray matter volume trajectories. **(B)** Quantification of individualized deviation scores (Z-scores) for each cerebellar lobule in SCA3 patients (red triangles) by projecting them onto the normative reference. **(C)** Identification of neuroanatomical biotypes using unsupervised clustering algorithm of deviation maps, followed by comparison of clinical characteristics. **(D)** Validation of biotype-specific therapeutic responses in a longitudinal subset of patients undergoing dTMS and comparative efficacy evaluation of dTMS versus rTMS. Abbreviations: GMV, gray matter volume; HCs, healthy controls; SCA3, spinocerebellar ataxia type 3; SARA, Scale for the Assessment and Rating of Ataxia; ICARS, International Cooperative Ataxia Rating Scale.

### Identification of Neurobiological Biotypes

To parse the neuroanatomical heterogeneity within SCA3, we applied a data-driven k-means clustering algorithm to the deviation profiles (Z-scores of 27 lobules) of the 114 patients (Figure 1C).To reduce the impact of random initial cluster centroids, we ran the clustering algorithm 100 times with distinct starting centers for each candidate cluster number.The number of clusters varied from 2 to 10, and the optimal number of clusters (*k*) was determined objectively using a consensus approach combining the silhouette coefficient and 24 validity indices from the NbClust package in R.^52^ This data-driven strategy ensures a reproducible classification independent of a priori hypotheses. Differences across biotypes were assessed with the two-sample t test or Mann–Whitney U test depending on data characteristics

### Clinical Predictive Modeling and Feature Importance

We investigated whether biotype-specific deviation patterns could predict clinical severity (SARA and ICARS scores).

#### Feature Selection

First, to reduce dimensionality, we calculated the effect size (Cohen’s *d*) of deviations between biotypes for each lobule. Lobules showing significant inter-biotype differences (FDR-corrected *q* < 0.05) were selected as feature vectors.

#### Ridge Regression Model

A ridge regression model with a nested 5-fold cross-validation (5F-CV) framework was employed. The inner loop optimized the penalty parameter, while the outer loop assessed generalization performance.^53,54^ Prediction accuracy was quantified using the pearson correlation coefficient (*r*) between predicted and observed scores. To ensure robustness, the 5F-CV process was repeated 101 times, reporting the median *r*. Statistical significance was assessed via 1,000 permutation tests.

#### Feature Weighting

The contribution of each lobule to symptom prediction was quantified by the ridge regression weight coefficients, where the absolute value indicates importance, and the sign indicates the direction of association.

### Validation of dTMS Treatment Response

To determine the clinical utility of the identified biotypes, we analyzed a longitudinal subset of 40 SCA3 patients who underwent a standardized dTMS intervention (Figure 1D).We employed linear mixed-effects models to evaluate the differential therapeutic effects of dTMS. A separate model was fitted for each clinical score (i.e., SARA and ICARS). The models included fixed effects of time point (pre-and post-intervention), Biotype (Biotype 1/Biotype 2), and the time point × Biotype interaction term; individual participant intercepts were modeled as random effects. To further quantify inter-biotype differences in therapeutic efficacy, Cohen’s d effect sizes were calculated. Treatment improvement was defined as the change (Δ) in SARA and ICARS score pre- and post-intervention. To clarify differences in therapeutic response to distinct TMS modalities across patients with different neuroanatomical biotypes, we used the neuroanatomical biotypes identified in this study as the stratification factor. Within each biotype, we compared changes in clinical score between rTMS and dTMS using the two-sample t test or the Mann–Whitney U test, where appropriate.

## Results

### Normative Modeling Unveils Substantial Neuroanatomical Heterogeneity

The normative model demonstrated excellent goodness-of-fit across all cerebellar lobules (Supplementary Fig. 2). When applied to the SCA3 cohort, the model revealed significant pathological deviations masked by group averages. Compared to healthy controls (HCs), SCA3 patients exhibited a significantly higher burden of structural deviations: both the total count of lobules with extreme deviations (*Cohen’s d* = 0.696, *p* < 0.001; Figure 2A) and the cumulative magnitude of deviations (*Cohen’s d*= 1.114, *p* < 0.001; Figure 2B) were markedly elevated.

**Figure 2.**
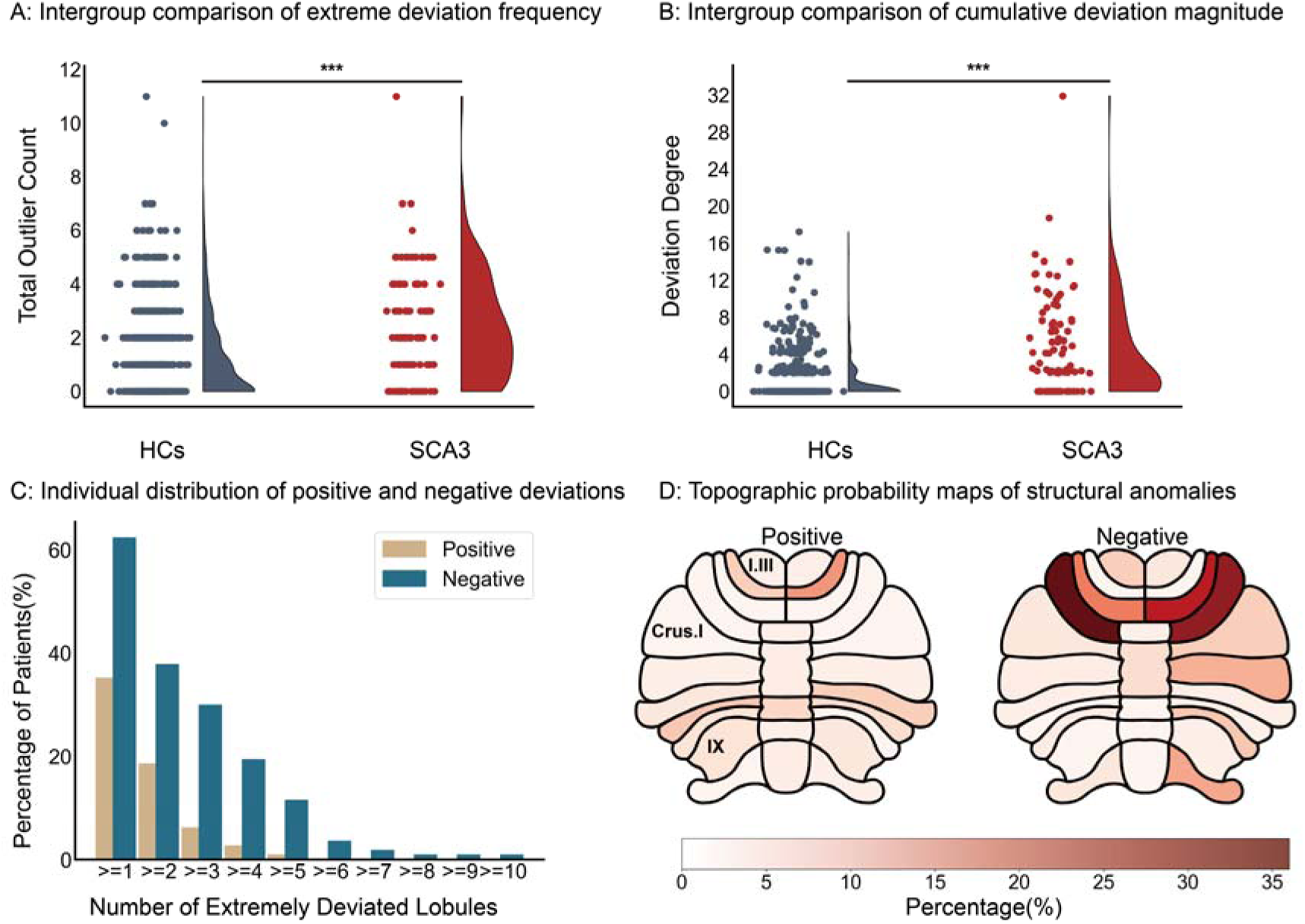
SCA3 patients exhibit significant individualized heterogeneity in cerebellar structural deviations. **(A, B)** Raincloud plots illustrating the increased burden of pathology in SCA3 compared to HCs. Patients show a significantly higher total count of outlier lobules **(A)** and a greater cumulative magnitude (degree) of deviations **(B)**. *** indicates *p* < 0.001. **(C)** Patient-level distribution of structural heterogeneity. The bar chart displays the count of lobules with extreme positive (beige) or negative (teal) deviations for each individual, highlighting diverse damage patterns. **(D)** Spatial overlap probability maps showing the frequency of extreme deviations across the cohort. The left panel maps the prevalence of positive deviations (structural preservation), while the right panel maps negative deviations (atrophy).

However, individual-level analysis highlighted profound heterogeneity. Notably, no single cerebellar lobule exhibited extreme deviations (*|Z*| > 1.96) in more than 35.96% of the patient cohort. For instance, while the Left Lobule VI showed the highest frequency of extreme deviations (35.96%), nearly 26.32% of patients showed no significant deviation in this region (Figure 2D). Nevertheless, over 62.28% of patients exhibited extreme deviations in at least one region (Figure 2C), confirming that SCA3 manifests as diverse, individualized neuroanatomical damage patterns rather than a uniform atrophy profile.

### Identification of Two Distinct Neurobiological Biotypes

A data-driven k-means clustering algorithm of the individual deviation maps identified two reproducible biotypes (*k*=2), a solution supported by the maximum silhouette coefficient and the highest voting frequency across all validity indices. (Figures 3A, 3B). The two biotypes displayed distinct, non-overlapping neuroanatomical profiles (Figure 3C):

**Figure 3.**
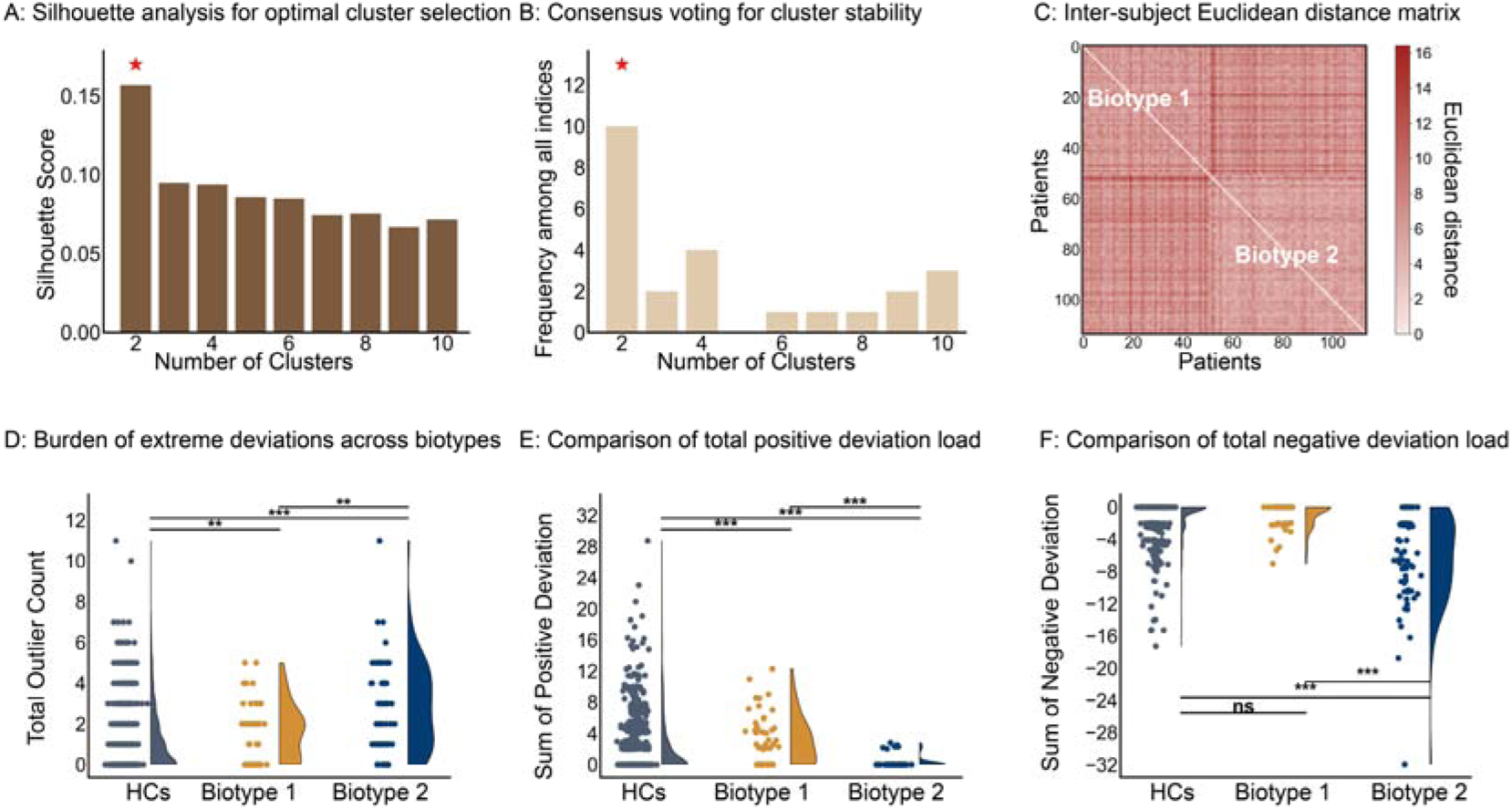
Identification of two distinct neuroanatomical biotypes using unsupervised clustering algorithm. **(A)** Silhouette analysis indicates that a cluster number of *k*=2 (red star) yields the highest coefficient, suggesting optimal separation. **(B)** Consensus voting by the NbClust package, where the majority of validity indices support a 2-cluster solution. **(C)** Similarity matrix based on Euclidean distance, visualizing the robust separation between Biotype 1 and Biotype 2. **(D–F)** Raincloud plots comparing deviation profiles. Biotype 2 is characterized by a significantly higher total outlier count **(D)** and extensive negative deviations **(F)**, whereas Biotype 1 is defined by predominant positive deviations **(E)**. Significance markers: * *p* < 0.05; ** *p* < 0.01; *** *p* < 0.001; ns, not significant.

#### Biotype 1 (Relative Preservation / Compensatory Profile, *n*=51)

Compared to HCs, this subgroup showed significantly more abnormal brain regions and a higher total positive deviation load (*Cohen’s d* = 0.302 and 0.320, respectively, *p* < 0.01; Figures 3D, 3E). Spatially, these “positive deviations” were predominantly characterized by structural integrity or potential compensatory hypertrophy in the posterior cerebellum (Bilateral VIIB, Bilateral VIIIA) and vermis (Posterior-Inferior regions), while sparing the anterior motor lobules (Figures 4A, 4C).

**Figure 4.**
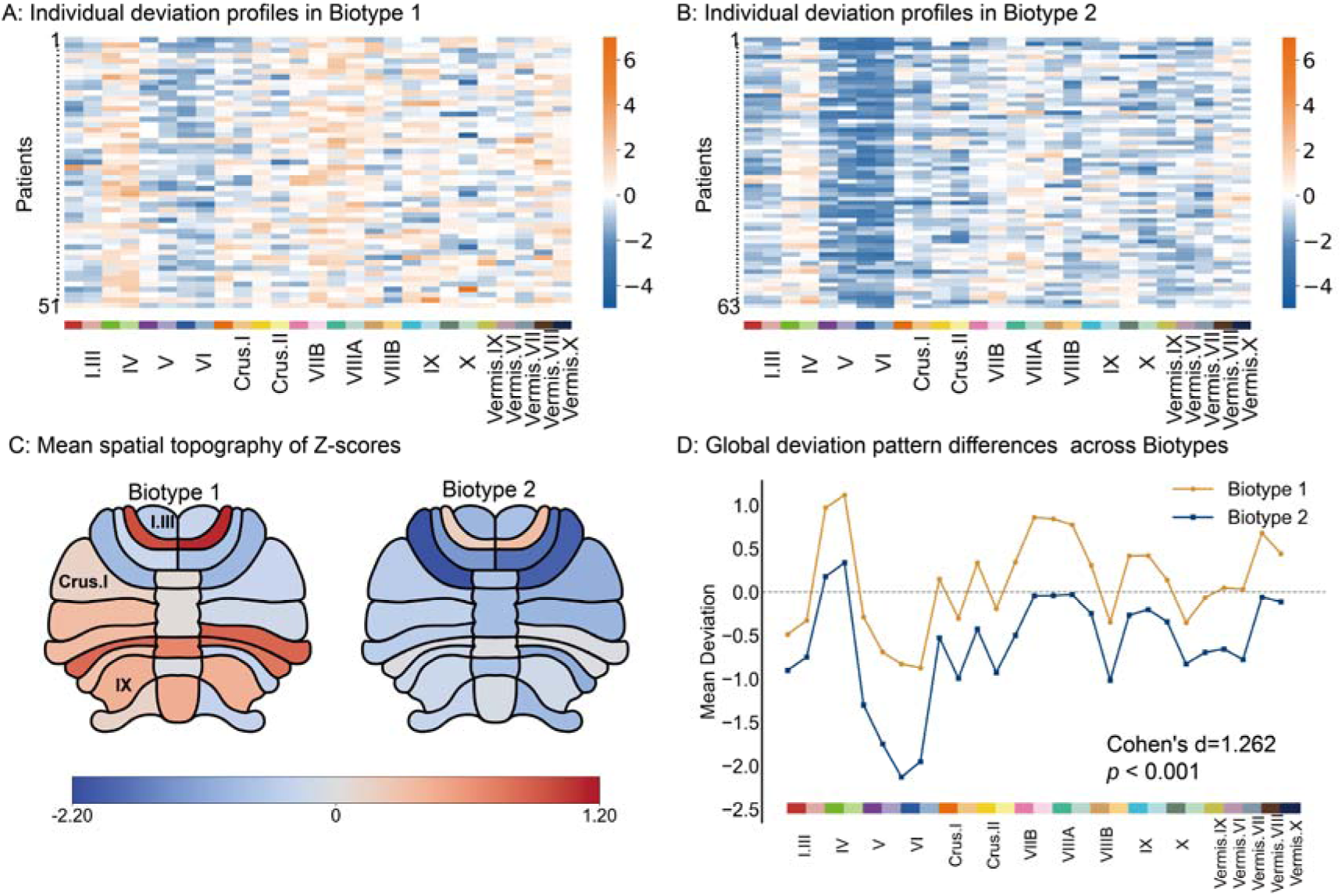
Divergent spatial patterns of cerebellar deviations between biotypes. (A, **B)** Individual deviation heatmaps for Biotype 1 **(A)** and Biotype 2 **(B)**. Columns represent 27 cerebellar lobules; rows represent patients. Blue indicates negative deviations (*Z* < 0, atrophy), and orange indicates positive deviations (*Z* > 0). **(C)** The mean deviation map of each biotype. Biotype 1 (left) shows relative preservation in the posterior cerebellum, while Biotype 2 (right) exhibits widespread anterior atrophy. **(D)** Global deviation pattern differences between biotypes. Biotype 2 (blue line) displays large negative deviations in anterior motor regions (Lobules I–VI), distinct from the profile of Biotype 1 (orange line).

#### Biotype 2 (Malignant Atrophy Profile, *n*=63)

In sharp contrast, Biotype 2 exhibited a “malignant” profile with significantly lower total positive deviations and negative deviations compared to HCs (*Cohen’s d* = -0.505 and -2.206, respectively, *p* < 0.001; Figures 3E, 3F). This biotype was defined by extensive, high-magnitude negative deviations concentrated in the anterior motor cerebellum (Bilateral I-VI) and extending to Crus I/II and the superior-middle vermis (Figures 4B, 4C).

The overall neuroimaging deviation patterns of the two biotypes were distinctly separated with a large effect size (*Cohen’s d* = 1.262, *p* < 0.001; Figure 4D).

### Anatomical Deviations Predict Clinical Severity: The Role of Right Lobule VIIB

To validate clinical relevance, A ridge regression model was performed using the 27 differentiating lobules as features (Figure 5A). In Biotype 2, deviation patterns significantly predicted clinical severity (SARA: *r* = 0.267, *p_perm_* = 0.019; ICARS: *r* = 0.237, *p_perm_* = 0.036; Figures 5B, 5C). In Biotype 1, no significant predictive association was found (SARA: *r* = 0.195, *p_perm_*= 0.111; ICARS: *r* = 0.143, *p_perm_* = 0.157).

**Figure 5.**
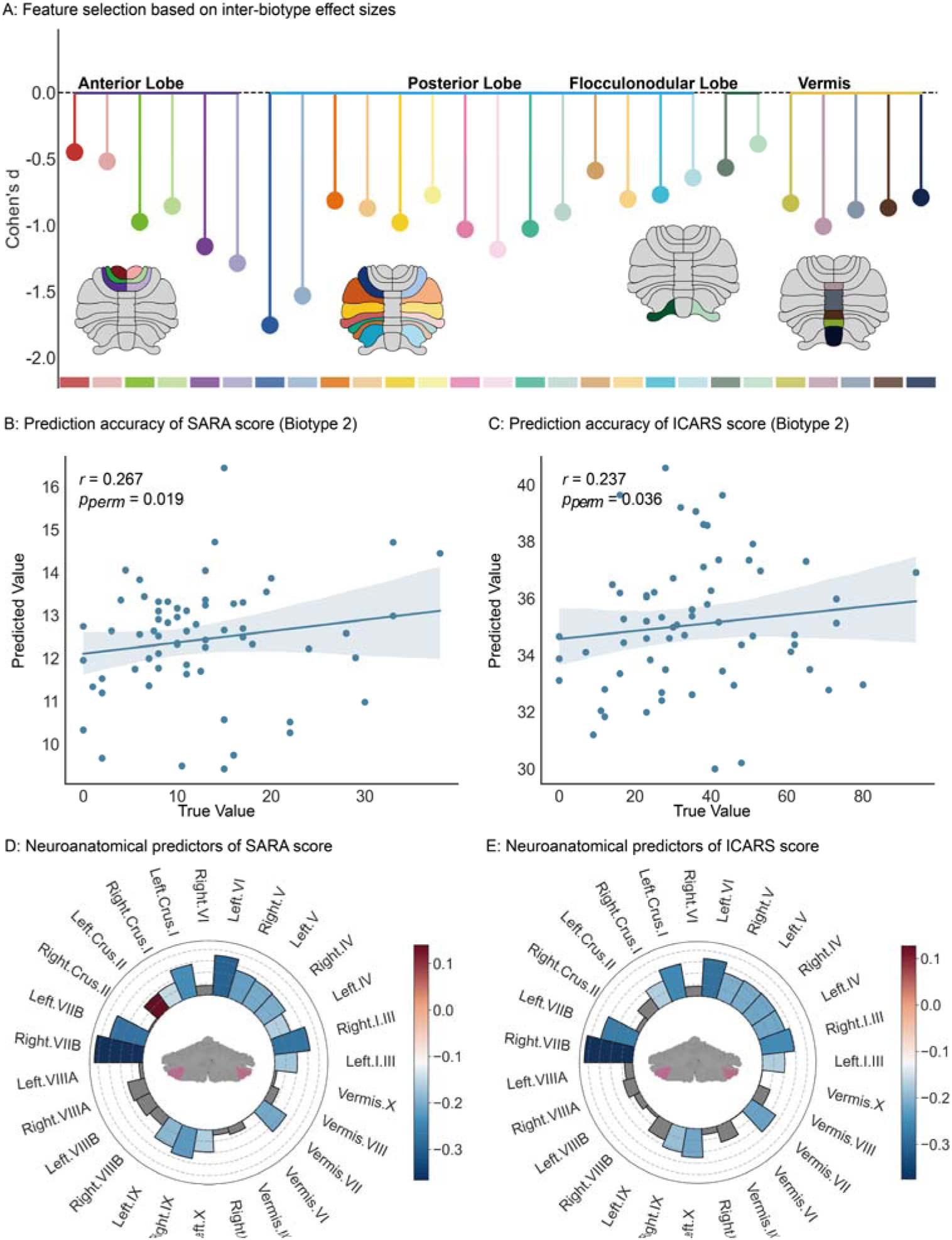
Biotype-specific structure-symptom coupling. **(A)** Lollipop chart showing lobules with significant inter-biotype differences (colored); non-significant lobules are in gray (*q* > 0.05). **(B, C)** Scatter plots of ridge regression model results in Biotype 2. Structural deviations significantly predict SARA score **(B)** and ICARS score **(C)**. The blue line represents the linear fit with 95% confidence intervals. **(D, E)** Feature weight distribution for SARA **(D)** and ICARS **(E)** prediction. Blue bars indicate negative weights (atrophy predicts severity). The Right Lobule VIIB shows the strongest negative contribution to clinical severity.

#### Feature Weight Analysis

The weights of significant features in the ridge regression model were predominantly negative, indicating that lower volume (atrophy) correlates with higher score (worse symptoms). Notably, the Right Lobule VIIB exhibited the largest absolute weight (Figures 5D, 5E), identifying it as the most critical neuroanatomical biomarker. This suggests that in the severe biotype, the structural collapse of the Right VIIB drives the worsening of ataxia.

### The “Symptom-Response Paradox”: Baseline Severity vs. dTMS Efficacy

Analysis of demographics revealed no significant differences in age, sex, and disease duration between biotypes (Table 2). However, they differed significantly in clinical status and therapeutic sensitivity:

**Table 2.** Comparative analysis of demographics, genetics, and clinical severity between SCA3 neuroanatomical biotypes.

| Characteristic | Biotype1(Compensatory)(n=51) | Biotype2(Atrophic)(n=63) | Statistic | P-value |
| --- | --- | --- | --- | --- |
| Demographics |  |  |  |  |
| Age, years | 40.18 ± 10.88 | 43.68 ± 11.83 | t = -1.63 | ns. |
| Sex, Male (n, %) | 27 (52.94%) | 37 (58.73%) | $\chi^2 = 0.18$ | ns. |
| Disease Characteristics |  |  |  |  |
| Disease Duration, years | 5.39 ± 4.93 | 6.56 ± 4.02 | U = 1264.5 | ns. |
| Age of Onset, years | 34.78 ± 11.10 | 37.13 ± 11.26 | U = 1383.5 | ns. |
| CAG Repeats | 69.98 ± 7.52 | 71.24 ± 6.32 | U = 1472.5 | ns. |
| Clinical Severity |  |  |  |  |
| SARA Score | 9.69 ± 7.33 | 12.5 ± 8.49 | U = 1306.0 | ns. |
| ICARS Score | 26.84 ± 19.19 | 35.44 ± 20.41 | U = 1197.0 | < 0.05 |

Footnotes: Values are expressed as mean ± SD or number (percentage). Note: Despite similar disease duration and genetic burden (CAG repeats), Biotype 2 exhibits significantly higher baseline clinical severity but greater therapeutic responsiveness. Abbreviations: SARA, Scale for the Assessment and Rating of Ataxia; ICARS, International Cooperative Ataxia Rating Scale.
The U values were obtained by the Mann–Whitney U test
The t value was obtained by the two-sample t test
The $\chi^2$ value for sex distribution was obtained by the Chi-square test
ns,not significant ( $p > 0.05$ )

#### Baseline Severity

Consistent with its atrophic profile, Biotype 2 presented with significantly more severe ataxia than Biotype 1 (SARA: *Cohen’s d* = 0.352, *p* =0.087; ICARS: *Cohen’s d* = 0.433, *p* < 0.05; Figures 6A, 6B).

**Figure 6.**
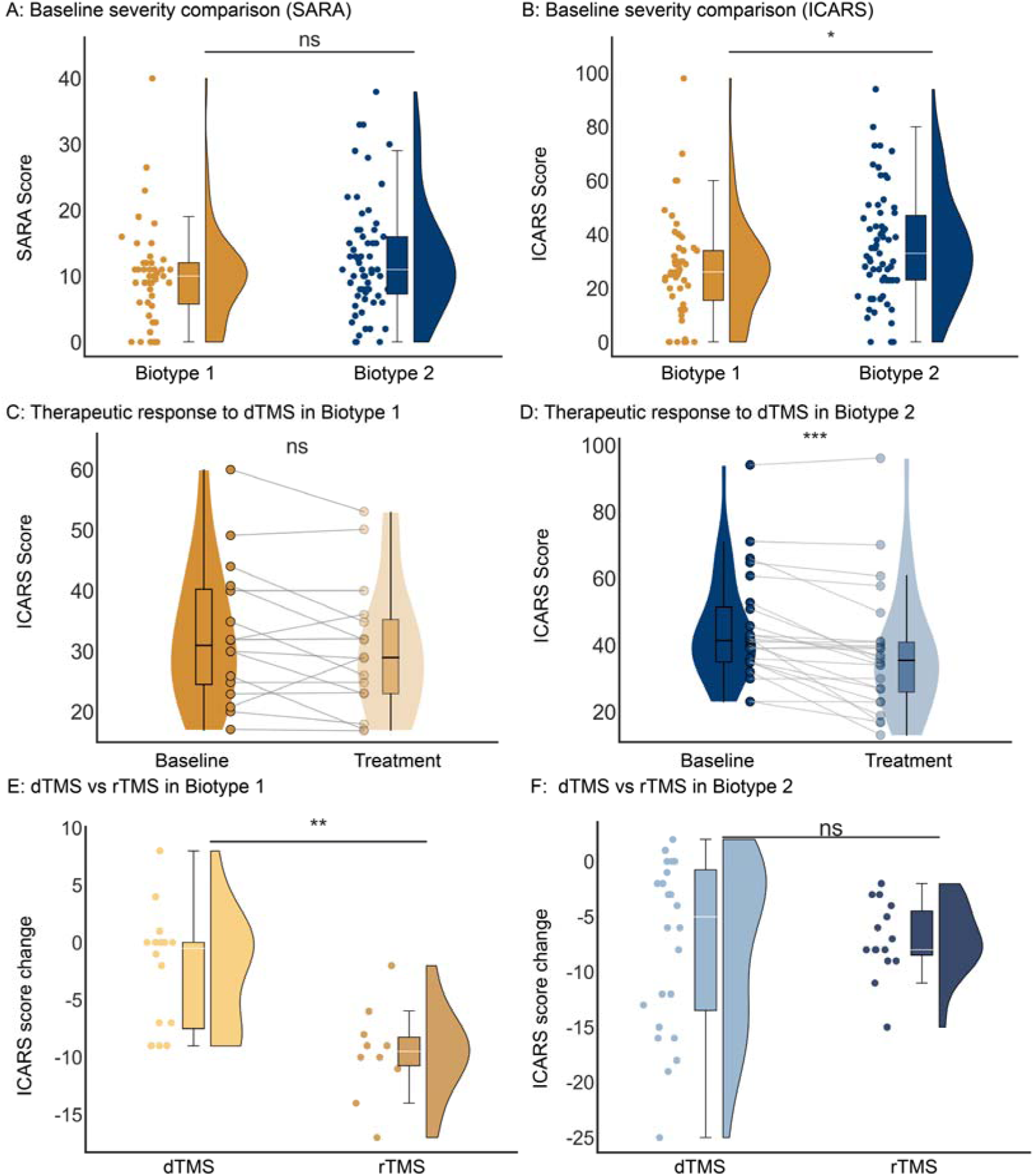
The “Symptom-Response Paradox”: Baseline severity vs. TMS efficacy. (A, **B)** Baseline clinical status. Biotype 2 patients exhibit significantly higher SARA **(A)** and ICARS **(B)** score compared to Biotype 1. **(C, D)** Treatment response. Despite worse baseline severity, Biotype 2 patients achieve greater therapeutic gains, reaching statistical significance for the ICARS scale **(D)**.(**E, F**) Therapeutic response difference: dTMS vs rTMS. Biotype 1 show greater ICARS score improvement with rTMS than with dTMS(**E**). Significance markers: * *p* < 0.05; ** *p* < 0.01; *** *p* < 0.001; ns, not significant. Boxplots represent the median (center line), interquartile range (box limits), and 1.5×IQR (whiskers).

#### dTMS Treatment Response

Analysis of the dTMS subset (Biotype 1, *n*=16; Biotype 2, *n*=24) revealed a striking “Symptom-Response Paradox.” Although Biotype 2 patients had worse baseline symptoms, they achieved superior therapeutic gains. While Biotype 1 showed no significant treatment effect (*p* > 0.05; Figure 6C), Biotype 2 manifested a significant reduction in symptoms (*p* < 0.001; Figure 6D). Notably, a significant time × biotype interaction (*p* = 0.026) was observed, statistically validating the differential treatment response across the two biotypes.The improvement in ICARS score after dTMS intervention was markedly greater in Biotype 2 than in Biotype 1, with a medium-to-large effect size (Cohen’s *d* = -0.719).A similar trend was observed for SARA score (*Cohen’s d* = - 0.436).These results demonstrate that patients with the most severe structural damage (Biotype 2) paradoxically exhibit the best clinical response to dTMS, particularly in motor coordination functions assessed by ICARS.

#### Comparison between rTMS and dTMS

Stratified according to the identified neuroanatomical biotypes, the rTMS treatment cohort included 10 patients classified as Biotype 1 and 15 as Biotype 2. Baseline characteristics including age, sex, SARA and ICARS scores were comparable between the dTMS and rTMS cohorts, with no statistically significant intergroup differences. For Biotype 1, the rTMS treatment resulted in significantly improvement in ataxia symptoms, with a larger reduction in ICARS score (*Cohen’s d* =-1.472, *p* < 0.01; Figure 6E) compared with dTMS. These findings indicated that rTMS could represent a promising treatment option for SCA3 patients classified as Biotype 1. In contrast, no significant between-group difference in therapeutic efficacy was observed for Biotype 2 (Figure 6F).

## Discussion

In this study, we addressed the longstanding challenge of clinical variability in spinocerebellar ataxia type 3 by constructing a disease-specific cerebellar normative model. Moving beyond the limitations of group-averaged atrophy maps, our approach enabled characterization of individualized neuroanatomical fingerprints across 114 patients. Our analysis identified two robust internally clustered neurobiological biotypes: a “Compensatory/Preserved” phenotype (Biotype 1), marked by preserved structural integrity within posterior cognitive cerebellar regions, and a “Malignant/Atrophic” phenotype (Biotype 2), featuring extensive degeneration of the anterior motor cerebellum. Most notably, longitudinal validation revealed an intriguing “Symptom-Response Paradox”: patients with severe structural degeneration (Biotype 2) displayed markedly better therapeutic gains following dTMS intervention. Collectively, these findings challenge the conventional “one-size-fits-all” paradigm and indicate that biological heterogeneity is not merely random noise but a promising candidate signal to inform future precision neuromodulation.

Clinical studies confirm marked phenotypic variability in SCA3, associated with unstable CAG repeat expansion and widespread central/peripheral neurodegeneration.^55–57^ Three classic clinical phenotypes, namely the Joseph type, Thomas type, and Machado type, are defined by onset age and dominant symptoms,^58^ but their classification is subject to subjective and environmental influences.^59^ Notably, small-sample longitudinal studies suggest MRI volumetric measures are more sensitive than clinical scales for tracking disease progression,^60–62^ supporting an objective neuroimaging-based approach to capturing SCA3 heterogeneity—consistent with our identification of neuroanatomical biotypes reflecting disease stages rather than CAG-driven biotypes.

The capacity to resolve such heterogeneity stems directly from the methodological shift offered by normative modeling.^63–65^ Unlike traditional case-control designs that mask inter-individual variability by focusing on mean differences, our approach accurately detected aberrant structural patterns at the individual level. Using this granular lens, we confirmed that while SCA3 patients globally exhibit widespread cerebellar atrophy, the spatial distribution of these anomalies is highly idiosyncratic. Notably, no single cerebellar lobule showed extreme deviations in more than 35.96% of the cohort, yet over 62.28% of patients manifested extreme alterations in at least one region. This result fundamentally challenges the view of SCA3 as a diffuse, uniform degeneration. Instead, it suggests that pathological heterogeneity is an intrinsic feature of the disease, necessitating the transition from group-based assumptions to the individualized deviation maps constructed here.

Leveraging this granular mapping, we identified Biotype 1 as a unique phenotype distinguished by “positive deviations,” predominantly in the posterior-inferior lobules (VIIB, VIIIA) and the vermis. We propose that these positive signals represent compensatory hypertrophy rather than mere structural preservation. According to the theory of cerebellar functional topography, the cerebellum possesses a “dual motor representation”: while the anterior lobe governs primary motor execution, lobules VIIB and VIIIA constitute secondary sensorimotor maps responsible for higher-order motor planning and sensory integration.^66,67^ In Biotype 1, the structural integrity—or potential expansion—of these posterior regions likely reflects a secondary circuit reorganization. As the anterior motor lobe begins to fail, these phylogenetically younger regions may undergo neuroplastic remodeling (e.g., increased dendritic arborization) to maintain functional homeostasis. This pattern provides direct imaging evidence for the “Cerebellar Reserve” hypothesis: the spared posterior cerebellum buffers the functional decline caused by incipient anterior pathology.^68^ Consequently, despite the disease presence, these patients maintain a relatively milder clinical phenotype.

In sharp contrast to this compensatory profile, Biotype 2 manifested a “malignant” neurodegenerative trajectory defined by extensive negative deviations. Anatomically, the profound atrophy in lobules V and VI—the primary motor representation areas—directly correlates with the functional collapse of sensorimotor integration, explaining the significantly more severe ataxia (higher SARA/ICARS score) observed in this subgroup^69^. Furthermore, the extension of pathology to Crus I and Crus II, key nodes of the default mode and executive control networks,^67^ suggests that Biotype 2 patients are at heightened risk for the Cerebellar Cognitive Affective Syndrome (CCAS). This identifies Biotype 2 not just as a motor phenotype, but as a severe, multi-system degeneration requiring comprehensive management. Further validating the clinical relevance of these anatomical distinctions, our ridge regression model analysis revealed a biotype-specific structure-symptom coupling. Specifically, feature weight analysis identified the Right Lobule VIIB as the most critical predictor with a dominant negative weight, indicating that the collapse of this specific region marks a tipping point in disease severity. Thus, the Right Lobule VIIB emerges as a sentinel biomarker for monitoring the transition from a compensated state to severe disability.

Perhaps the most clinically impactful finding, however, lies in the dissociation between baseline severity and therapeutic potential—a phenomenon we term the “Symptom-Response Paradox.” Although Biotype 2 patients presented with greater structural damage and symptom burden, they paradoxically achieved superior therapeutic gains from dTMS compared to Biotype 1. We interpret this seemingly counterintuitive finding through the lens of state-dependent neuroplasticity and homeostatic metaplasticity. Since the neural circuits in Biotype 1 are likely already operating at near-maximal compensatory capacity (as evidenced by positive deviations), their potential for further facilitation by exogenous dTMS is saturated (“ceiling effect”). Conversely, the atrophied motor cortex in Biotype 2 likely exists in a state of hypoactivity or excessive inhibition. According to the Bienenstock-Cooper-Munro (BCM) theory, neural networks in a suppressed state have a lower threshold for long-term potentiation (LTP) and a larger dynamic range for synaptic strengthening.^70^ Therefore, dTMS may exert its maximal leverage by rescuing these “silenced” residual circuits in Biotype 2.

Furthermore, we identified biotype-specific differences in therapeutic responses to distinct TMS modalities across neuroanatomical biotypes. These findings provide an evidence-based rationale to tackle the longstanding clinical challenge of TMS regimen selection for patients with spinocerebellar ataxia type 3 (SCA3). To date, no biotype-specific evidence-based clinical guidelines are available to guide selection between conventional rTMS and dTMS for SCA3, and treatment decisions remain largely empirical and unstandardized in routine clinical practice. Our comparative analysis partially addresses this important evidence gap: among Biotype 1 patients, conventional rTMS was associated with significantly larger reductions in ICARS score and greater symptomatic improvement relative to dTMS. This result offers preliminary evidence supporting preferential rTMS use in this patient subgroup.

These observations carry direct translational implications for future clinical trial design. Variable treatment responses commonly reported in ataxia trials may partly arise from underrecognized mixing of robust responders (Biotype 2) and poor responders (Biotype 1) within unstratified cohorts. We propose that future trials move beyond uniform “one-size-fits-all” enrollment criteria. Instead, normative modeling-derived Z-scores could act as stratifying biomarkers: individuals with a Biotype 2-like anatomical profile (high negative deviation load in anterior motor lobules) may be suitable candidates for excitability-enhancing stimulation protocols, while patients presenting with the compensatory Biotype 1 phenotype could be considered for prioritized enrollment in rTMS-focused trials, given their robust symptomatic improvements following rTMS intervention.

A critical issue concerning neuroimaging biotypes identified through cerebellar normative modeling is whether these biotypes merely represent intuitive reflections of different stages of disease progression. The data from the present study showed no statistically significant difference in disease duration between the two identified biotypes, confirming that these biotypes reflect intrinsic biological differences among patients rather than simply corresponding to distinct stages of disease development. Another key issue to be investigated is whether the differences among SCA3 biotypes can be explained by the length of the CAG repeat, the primary genetic driver of this disease. The results of the present study showed no statistically significant differences in CAG repeat length among the identified biotypes. Our findings are consistent with previous studies: CAG repeat length only accounts for approximately 50%–70% of the variance in age at onset, and cannot reliably predict disease progression rate or cerebellar functional decline.^71–73^ Phenotypic discordance has been reported in monozygotic twins with SCA3, further supporting the limited explanatory power of CAG repeat length for disease phenotypes.^74^The two biotypes identified by neuroimaging in the present study may serve as specific endophenotypes of SCA3, providing a promising and practical framework for precise patient stratification and individualized therapy in future clinical research.

Several limitations warrant consideration in interpreting these results. First, while our normative model was trained on a large multi-center cohort, the clinical validation in SCA3 was performed at a single center. Multi-center replication is needed to ensure generalizability. Second, our clustering relied solely on gray matter volume; integrating diffusion tensor imaging based tractography and functional connectivity could offer a more holistic view of the “compensatory” versus “degenerative” networks. Third, our analysis of biotype-specific therapeutic responses to rTMS and dTMS was derived from a longitudinal interventional subset of the cohort with a relatively small sample size. Future randomized, sham-controlled trials stratified by these identified neuroanatomical biotypes are essential to validate both the durability of the observed treatment effects and the predictive utility of our biotype classification for personalized treatment selection. Finally, longitudinal tracking of individual Z-score trajectories is required to determine whether Biotype 1 eventually collapses into Biotype 2, describing a continuum of disease progression.

In conclusion, this study systematically delineated the core characteristics of cerebellar neuroanatomical heterogeneity in SCA3 using a normative modeling framework. We identified two neurobiological biotypes that reflect distinct pathophysiological stages, i.e., “Compensatory” versus “Decompensated”. Crucially, we revealed that these biotypes possess divergent sensitivities to neuromodulation. These findings not only underscore the necessity of individualized neuroanatomical assessment but also provide an actionable, imaging-based roadmap for precision medicine, ensuring that the right patient receives the optimal intervention strategy.

## Data availability

The Human Connectome Project-Young Adult and Human Connectome Project-Aging datasets are publicly available at https://www.humanconnectome.org/. The Alzheimer’s Disease Neuroimaging Initiative dataset is freely accessible at http://adni.loni.usc.edu/. All remaining datasets generated or analyzed during the current study will be made available from the corresponding author upon reasonable request.

## Code availability

The neuroimaging preprocessing in this study was performed using the ACAPULCO pipeline, which is publicly accessible at (https://gitlab.com/shuohan/acapulco). Furthermore, the normative modeling was implemented via the Predictive Clinical Neuroscience toolkit, an open-source Python package available at (https://github.com/amarquand/PCNtoolkit).

## ACKNOWLEDGMENTS AND DISCLOSURES

This work was partially supported by Chongqing Science and Health Joint Medical Research Key Project (Grant No. 2025GGXM005), the National Natural Science Foundation of China (Grant Nos. 82472061), the Beijing Natural Science Foundation (Grant No.7262231) and the Yantai “Double Hundred Plan” Talent Grant. Our heartfelt thanks are also due to all participating patients and their families for their selfless dedication of time, dedication, and enthusiastic cooperation.

## Author contributions

K. K. W. performed the experimental studies; K. K. W. and L. Z. F. mainly wrote the manuscript; Y. W. H. and X. G. W. performed the collection and collation of the clinical data; C. Y. C., L. Z. F., and C. L. provided theoretical guidance and informed interpretation of the results; K. K. W., L. Z. F.,and C. L. reviewed the manuscript. L. Z. F., and C. L. supervised the project. All authors discussed the results and commented on the manuscript at all stages.

## References

1. Klockgether T, Mariotti C, Paulson HL. Spinocerebellar ataxia. Nat Rev Dis Primers.2019;5(1):24.

2. Riess O, Rub U, Pastore A, Bauer P, Schols L. SCA3: neurological features, pathogenesis and animal models. Cerebellum. 2008;7(2):125–137.

3. Vaz RL, Ribeiro GR, Nery LG, da Costa ACMM, Oliveira GS, Arruda JT. Neuropatologia da ataxia espinocerebelar tipo 3 (SCA3)–a doença de Machado-Joseph. Research, Society and Development. 2021;10(3):e16910313138–e16910313138.

4. Bostan AC, Strick PL. The basal ganglia and the cerebellum: nodes in an integrated network. Nat Rev Neurosci. 2018;19(6):338–350.

5. Ribeiro RFN, Pereira D, Lopes SM, et al. Circadian rhythms are disrupted in patients and preclinical models of Machado-Joseph disease. Brain. 2025;148(11):4127–4142.

6. Brooker SM, Edamakanti CR, Akasha SM, Kuo SH, Opal P. Spinocerebellar ataxia clinical trials: opportunities and challenges. Ann Clin Transl Neurol. 2021;8(7):1543–1556.

7. Silva P, Costa MA, Gaspar L, et al. The Medication Patterns of Spinocerebellar Ataxia Type 3 Mutation Carriers Enrolled in the ESMI Cohort. CNS Drugs.2025.

8. Shi Y, Zou G, Chen Z, et al. Efficacy of cerebellar transcranial magnetic stimulation in spinocerebellar ataxia type 3: a randomized, single-blinded, controlled trial. J Neurol.2023;270(11):5372–5379.

9. Sikandar A, Liu XH, Xu HL, et al. Short-term efficacy of repetitive transcranial magnetic stimulation in SCA3: A prospective, randomized, double-blind, sham-controlled study. Parkinsonism Relat Disord. 2023;106:105236.

10. Franca C, de Andrade DC, Silva V, et al. Effects of cerebellar transcranial magnetic stimulation on ataxias: A randomized trial. Parkinsonism Relat Disord. 2020;80:1–6.

11. Liu X, Guo J, Wang X, et al. Regional brain atrophy subtypes in spinocerebellar ataxia type 3: links to clinical performance and treatment response. J Neurol. 2025;273(1):44.

12. Guo J, Chen H, Biswal BB, et al. Gray matter atrophy patterns within the cerebellum-neostriatum-cortical network in SCA3. Neurology. 2020;95(22):e3036–e3044.

13. Peng H, Liang X, Long Z, et al. Gene-Related Cerebellar Neurodegeneration in SCA3/MJD: A Case-Controlled Imaging-Genetic Study. Front Neurol. 2019;10:1025.

14. Camargos ST, Marques W, Jr., Santos AC. Brain stem and cerebellum volumetric analysis of Machado Joseph disease patients. Arq Neuropsiquiatr. 2011;69(2B):292–296.

15. Shi L, Ou L, Ou P, et al. Gray Matter Asymmetry Alterations in Patients With Spinocerebellar Ataxia Type 3: A Voxel-Based Morphometric Comparison Study. CNS Neurosci Ther. 2024;30(12):e70171.

16. D’Abreu A, Franca MC, Jr., Yasuda CL, Campos BA, Lopes-Cendes I, Cendes F. Neocortical atrophy in Machado-Joseph disease: a longitudinal neuroimaging study. J Neuroimaging. 2012;22(3):285–291.

17. Ogawa Y, Ito S, Makino T, Kanai K, Arai K, Kuwabara S. Flattened facial colliculus on magnetic resonance imaging in Machado-Joseph disease. Mov Disord.2012;27(8):1041–1046.

18. Sobana SA, Huda F, Hermawan R, et al. Brain MRI Volumetry Analysis in an Indonesian Family of SCA 3 Patients: A Case-Based Study. Front Neurol. 2022;13:912592.

19. Ye ZX, Bi J, Qiu LL, et al. Cognitive impairment associated with cerebellar volume loss in spinocerebellar ataxia type 3. J Neurol. 2024;271(2):918–928.

20. Stefanescu MR, Dohnalek M, Maderwald S, et al. Structural and functional MRI abnormalities of cerebellar cortex and nuclei in SCA3, SCA6 and Friedreich’s ataxia. Brain. 015;138(Pt 5):1182–1197.

21. Petit E, Coarelli G, Morgan D, et al. Predictive models for ataxia progression and conversion in spinocerebellar ataxia type 1 and 3. Brain. 2025.

22. Hu J, Chen X, Li M, et al. Pattern of cerebellar grey matter loss associated with ataxia severity in spinocerebellar ataxias type 3: a multi-voxel pattern analysis. Brain Imaging Behav. 2022;16(1):379–388.

23. Qiu H, Wu C, Liang J, et al. Structural alterations of spinocerebellar ataxias type 3: from pre-symptomatic to symptomatic stage. Eur Radiol. 023;33(4):2881–2894.

24. Marquand AF, Rezek I, Buitelaar J, Beckmann CF. Understanding Heterogeneity in Clinical Cohorts Using Normative Models: Beyond Case-Control Studies. Biol Psychiatry. 016;80(7):552–561.

25. Marquand AF, Kia SM, Zabihi M, Wolfers T, Buitelaar JK, Beckmann CF. Conceptualizing mental disorders as deviations from normative functioning. Mol Psychiatry. 2019;24(10):1415–1424.

26. Marquand AF, Wolfers T, Mennes M, Buitelaar J, Beckmann CF. Beyond Lumping and Splitting: A Review of Computational Approaches for Stratifying Psychiatric Disorders. Biol Psychiatry Cogn Neurosci Neuroimaging. 2016;1(5):433–447.

27. Bethlehem RAI, Seidlitz J, White SR, et al. Brain charts for the human lifespan. Nature. 2022;604(7906):525–533.

28. Sun L, Zhao T, Liang X, et al. Human lifespan changes in the brain’s functional connectome. Nat Neurosci. 2025;28(4):891–901.

29. Cole TJ. The development of growth references and growth charts. Ann Hum Biol. 2012;39(5):382–394.

30. Sun X, Sun J, Lu X, et al. Mapping Neurophysiological Subtypes of Major Depressive Disorder Using Normative Models of the Functional Connectome. Biol Psychiatry.2023;94(12):936–947.

31. Shan X, Uddin LQ, Xiao J, et al. Mapping the Heterogeneous Brain Structural Phenotype of Autism Spectrum Disorder Using the Normative Model. Biol Psychiatry. 2022;91(11):967–976.

32. Bu X, Zhao Y, Zheng X, et al. Normative growth modeling of brain morphology reveals neuroanatomical heterogeneity and biological subtypes in children with ADHD. *bioRxiv*. 2024:2024.2003. 2016.582202.

33. Kim M, Leonardsen E, Rutherford S, et al. Mapping cerebellar anatomical heterogeneity in mental and neurological illnesses. Nature Mental Health. 2024;2(10):1196–1207.

34. Gaiser C, van der Vliet R, de Boer AAA, et al. Population-wide cerebellar growth models of children and adolescents. Nat Commun. 2024;15(1):2351.

35. Kim M, Sharma N, Leonardsen EH, et al. Predicting Mental and Neurological Illnesses Based on Cerebellar Normative Features. Biol Psychiatry Glob Open Sci. 2025;5(5):100541.

36. Van Essen DC, Smith SM, Barch DM, et al. The WU-Minn Human Connectome Project: an overview. Neuroimage.2013;80:62–79.

37. Bookheimer SY, Salat DH, Terpstra M, et al. The Lifespan Human Connectome Project in Aging: An overview. Neuroimage. 2019;185:335–348.

38. Jack CR, Jr., Bernstein MA, Fox NC, et al. The Alzheimer’s Disease Neuroimaging Initiative (ADNI): MRI methods. J Magn Reson Imaging. 2008;27(4):685–691.

39. Jack CR, Jr., Bennett DA, Blennow K, et al. NIA-AA Research Framework: Toward a biological definition of Alzheimer’s disease. Alzheimers Dement. 2018;14(4):535–562.

40. Trouillas P, Takayanagi T, Hallett M, et al. International Cooperative Ataxia Rating Scale for pharmacological assessment of the cerebellar syndrome. The Ataxia Neuropharmacology Committee of the World Federation of Neurology. J Neurol Sci. 1997;145(2):205–211.

41. Schmitz-Hubsch T, du Montcel ST, Baliko L, et al. Scale for the assessment and rating of ataxia: development of a new clinical scale. Neurology. 2006;66(11):1717–1720.

42. Han S, An Y, Carass A, Prince JL, Resnick SM. Longitudinal analysis of regional cerebellum volumes during normal aging. Neuroimage. 2020;220:117062.

43. Carass A, Cuzzocreo JL, Han S, et al. Comparing fully automated state-of-the-art cerebellum parcellation from magnetic resonance images. Neuroimage. 2018;183:150–172.

44. Tustison NJ, Avants BB, Cook PA, et al. N4ITK: improved N3 bias correction. IEEE Trans Med Imaging. 2010;29(6):1310–1320.

45. Fonov V, Evans AC, Botteron K, et al. Unbiased average age-appropriate atlases for pediatric studies. Neuroimage. 2011;54(1):313–327.

46. Rutherford S, Kia SM, Wolfers T, et al. The normative modeling framework for computational psychiatry. Nat Protoc. 2022;17(7):1711–1734.

47. Bayer JMM, van Velzen LS, Pozzi E, et al. Dissecting heterogeneity in cortical thickness abnormalities in major depressive disorder: a large-scale ENIGMA MDD normative modelling study. *bioRxiv*.2025.

48. Fraza CJ, Dinga R, Beckmann CF, Marquand AF. Warped Bayesian linear regression for normative modelling of big data. Neuroimage.2021;245:118715.

49. Kia SM, Huijsdens H, Rutherford S, et al. Federated multi-site normative modeling using hierarchical Bayesian regression. BioRxiv. 2021:2021.2005. 2028.446120.

50. Bayer JMM, Dinga R, Kia SM, et al. Accommodating site variation in neuroimaging data using normative and hierarchical Bayesian models. Neuroimage. 2022;264:119699.

51. Dinga R, Fraza CJ, Bayer JM, Kia SM, Beckmann CF, Marquand AF. Normative modeling of neuroimaging data using generalized additive models of location scale and shape. BioRxiv. 2021:2021.2006. 2014.448106.

52. Charrad M, Ghazzali N, Boiteau V, Niknafs A. NbClust: an R package for determining the relevant number of clusters in a data set. Journal of statistical software. 2014;61:1–36.

53. Cui Z, Pines AR, Larsen B, et al. Linking Individual Differences in Personalized Functional Network Topography to Psychopathology in Youth. Biol Psychiatry. 2022;92(12):973–983.

54. Cui Z, Li H, Xia CH, et al. Individual Variation in Functional Topography of Association Networks in Youth. Neuron. 2020;106(2):340–353 e348.

55. Roy SK, Liu X (2021) Emerging concepts of pathogenesis and comprehensive therapeutic strategies for spinocerebellar ataxia type 3. Neurosci Med 12(01):22–43

56. Schols L, Bauer P, Schmidt T, Schulte T, Riess O. Autosomal dominant cerebellar ataxias: clinical features, genetics, and pathogenesis. Lancet Neurol.2004;3(5):291–304.

57. Koeppen AH. The Neuropathology of Spinocerebellar Ataxia Type 3/Machado-Joseph Disease. Adv Exp Med Biol. 2018;1049:233–241.

58. Coutinho P, Andrade C. Autosomal dominant system degeneration in Portuguese families of the Azores Islands. A new genetic disorder involving cerebellar, pyramidal, extrapyramidal and spinal cord motor functions. Neurology. 1978;28(7):703–709.

59. Da Silva JD, Teixeira-Castro A, Maciel P. From Pathogenesis to Novel Therapeutics for Spinocerebellar Ataxia Type 3: Evading Potholes on the Way to Translation. Neurotherapeutics. 2019;16(4):1009–1031.

60. Piccinin CC, Rezende TJR, de Paiva JLR, et al. A 5-Year Longitudinal Clinical and Magnetic Resonance Imaging Study in Spinocerebellar Ataxia Type 3. Mov Disord. 2020;35(9):1679–1684.

61. Adanyeguh IM, Perlbarg V, Henry PG, et al. Autosomal dominant cerebellar ataxias: Imaging biomarkers with high effect sizes. Neuroimage Clin. 2018;19:858–867.

62. Reetz K, Rodriguez-Labrada R, Dogan I, et al. Brain atrophy measures in preclinical and manifest spinocerebellar ataxia type 2. Ann Clin Transl Neurol. 2018;5(2):128–137.

63. Verdi S, Kia SM, Yong KXX, et al. Revealing Individual Neuroanatomical Heterogeneity in Alzheimer Disease Using Neuroanatomical Normative Modeling. Neurology.2023;100(24):e2442–e2453.

64. Verdi S, Rutherford S, Fraza C, et al. Personalizing progressive changes to brain structure in Alzheimer’s disease using normative modeling. Alzheimers Dement. 2024;20(10):6998–7012.

65. Fraza C, Buckova BR, Johansson ME, Helmich RC, Marquand AF, Beckmann CF. Understanding individual neurodegenerative progression in Parkinson’s disease through normative modelling. Sci Rep. 2025;15(1):36659.

66. Stoodley CJ, Schmahmann JD. Functional topography in the human cerebellum: a meta-analysis of neuroimaging studies. Neuroimage. 2009;44(2):489–501.

67. Buckner RL, Krienen FM, Castellanos A, Diaz JC, Yeo BT. The organization of the human cerebellum estimated by intrinsic functional connectivity. J Neurophysiol. 2011;106(5):2322–2345.

68. Mitoma H, Buffo A, Gelfo F, et al. Consensus Paper. Cerebellar Reserve: From Cerebellar Physiology to Cerebellar Disorders. Cerebellum. 2020;19(1):131–153.

69. King M, Hernandez-Castillo CR, Poldrack RA, Ivry RB, Diedrichsen J. Functional boundaries in the human cerebellum revealed by a multi-domain task battery. Nat Neurosci. 2019;22(8):1371–1378.

70. Bienenstock EL, Cooper LN, Munro PW. Theory for the development of neuron selectivity: orientation specificity and binocular interaction in visual cortex. J Neurosci. 1982;2(1):32–48.

71. Matilla-Duenas A, Ashizawa T, Brice A, et al. Consensus paper: pathological mechanisms underlying neurodegeneration in spinocerebellar ataxias. Cerebellum. 2014;13(2):269–302.

72. Tezenas du Montcel S, Durr A, Rakowicz M, et al. Prediction of the age at onset in spinocerebellar ataxia type 1, 2, 3 and 6. J Med Genet. 2014;51(7):479–486.

73. Huang SR, Wu YT, Jao CW, et al. CAG repeat length does not associate with the rate of cerebellar degeneration in spinocerebellar ataxia type 3. Neuroimage Clin. 2017;13:97–105.

74. Zhao H, Yang L, Dong Y, Wu ZY. Phenotypic variance in monozygotic twins with SCA3. Mol Genet Genomic Med. 2020;8(10):e1438.

